# Constraint Network Analysis of Global and Local Rigidity in Wild-Type EGFR: Apo vs Gefitinib-Bound States

**DOI:** 10.64898/2026.01.19.700489

**Authors:** Kankana Bhattacharjee, Aryya Ghosh

## Abstract

The mechanical rigidity–flexibility architecture of protein kinases play a critical role in regulating conformational stability and inhibitor response, yet remains insufficiently quantified for Epidermal Growth Factor Receptor (EGFR). Here, we apply Constraint Network Analysis (CNA) to systematically characterize the global and local mechanical properties of wild-type EGFR in its apo state and when bound to the ATP-competitive inhibitor gefitinib. Analysis of multiple global rigidity indices reveal a well-defined rigidity percolation transition in apo EGFR at an energy cutoff of approximately −2.0 kcal mol^−1^, indicative of an intrinsically stable mechanical framework. Gefitinib binding shifts this transition slightly to higher energies and sharpens the percolation behavior, accompanied by enhanced long-range mechanical coupling, increased rigidity order parameters, and reduced configurational entropy. Importantly, residue-level rigidity and percolation profiles remain largely conserved between the two states, demonstrating that ligand binding does not induce large-scale reorganization of the EGFR mechanical network. Instead, inhibition arises from subtle, localized rigidification within functionally relevant regions, consistent with stabilization of an inactive conformational ensemble. Collectively, this work establishes the first CNA-based mechanical reference state for wild-type EGFR and underscores the utility of network rigidity analysis for resolving ligand-induced effects that are structurally subtle yet mechanistically significant. This framework provides a quantitative baseline for future studies of oncogenic mutations and drug-resistant EGFR variants.

## Introduction

Protein rigidity and flexibility play a central role in determining structural stability, catalytic efficiency, and molecular recognition. Proteins are not mechanically uniform; instead, they display heterogeneous distributions of rigid and flexible regions, which enable the conformational adaptability required for biological function. These mechanical features are often evolutionarily conserved among homologous proteins and can be fine-tuned across organisms to accommodate physiological temperature and environmental conditions. Consequently, identifying mechanically distinct regions within proteins and understanding their responses to perturbations such as ligand binding, thermal variations, or changes in solvent environment are essential for elucidating functional mechanisms and for advancing rational protein engineering and structure-based drug design strategies [1–8].

A variety of experimental techniques, including hydrogen–deuterium exchange, neutron scattering, and NMR relaxation experiments, have provided valuable insights into biomolecular flexibility. Despite these advances, fully capturing the breadth of protein motions across relevant spatial and temporal scales remains challenging. In many cases, functionally important dynamics arise from subtle, localized fluctuations that do not produce large conformational changes detectable by conventional structural analyses. Computational approaches therefore offer a critical complementary perspective by enabling systematic and scalable characterization of protein rigidity and flexibility at atomic resolution [9–11].

From a broader perspective, protein function can be understood as an emergent property of network organization operating across multiple scales. At the systems level, protein–protein interaction (PPI) networks exhibit characteristic topological features such as scale-free connectivity, hierarchical modularity, and fractal organization, which underpin robustness, information flow, and functional specialization. Network-theoretical analyses of disease-associated PPI networks have demonstrated that perturbations often propagate through well-defined regulatory backbones rather than uniformly across the network, highlighting the importance of structurally and topologically central nodes [12, 13]. Constraint Network Analysis extends this network-based viewpoint to the intramolecular scale by representing proteins themselves as mechanical networks of constraints. In this framework, rigidity and flexibility emerge from the topology and strength of interactions within the network, providing a natural bridge between systems-level network organisation and atomic-level mechanical stability. Constraint Network Analysis (CNA), as implemented in the FIRST software, represents biomolecules as networks of covalent and noncovalent constraints and employs the Pebble Game algorithm to distinguish rigid clusters from flexible regions. This physics-based framework allows quantitative assessment of mechanical stability without the need for extensive conformational sampling. CNA has been successfully applied to a wide range of biomolecular systems to investigate conformational stability, protein–protein interactions, and the mechanical underpinnings of allosteric regulation [14–19]. Dasetty *et al*. showed that alterations in residue–residue interaction networks, driven by subtle changes in flexibility and long-range correlations, can critically influence protein function without inducing large-scale structural rearrangements. These findings underscore the utility of network-based approaches for resolving mechanistically relevant but structurally subtle effects, providing strong motivation for applying constraint-based rigidity analysis to kinase systems [20].

Although the Epidermal Growth Factor Receptor (EGFR) has been extensively characterized using structural biology, molecular dynamics simulations, and mutational studies, its intrinsic rigidity–flexibility organization in the context of ligand binding remains incompletely understood [21]. Protein kinases, in particular, are known to possess highly conserved mechanical frameworks that support catalytic activity, which can obscure subtle allosteric effects when analyzed using traditional structural descriptors [22]. Notably, although gefitinib binding induces a modest shift and sharpening of the global rigidity percolation transition, the overall global and residue-level rigidity–flexibility profiles of wild-type EGFR remain highly conserved. This behavior indicates that the kinase preserves its intrinsic mechanical architecture upon inhibitor engagement, with ligand binding primarily enhancing mechanical coupling and local stabilization rather than driving a qualitative reorganization of the rigidity– flexibility network.

In this work, we present the first systematic CNA-based analysis of EGFR–Gefitinib interactions. By integrating network rigidity metrics with established structural features of EGFR, this study provides a mechanistic framework for understanding inhibitor tolerance and establishes a foundation for future comparative analyses involving oncogenic or drug-resistant EGFR variants.

### Computational Methodology

Due to the absence of experimentally resolved crystal structure for wild-type EGFR in complex with Gefitinib, molecular docking was employed to generate reliable initial binding conformation for these inhibitor. To resolve this, the initial structure for the wild-type EGFR– Erlotinib complex was obtained from the RCSB Protein Data Bank (PDB ID: 1M17), which provides a high-resolution X-ray crystallographic structure suitable for molecular modelling.

Docking calculation was carried out using the GOLD 2020.3 software package [23], which applies a genetic algorithm-based search strategy to explore ligand binding poses within the defined active site. The crystal structure of EGFR bound to Erlotinib (PDB ID: 1M17) was utilized as a structural template to ensure consistency in receptor conformation across docking study.

For the ligand i.e, Gefitinib, multiple docking poses were generated and ranked according to the GOLD fitness score, a scoring function that evaluates ligand–protein complementarity based on hydrogen bonding, van der Waals interactions, and ligand strain. The top-scoring pose for ligand corresponding to the highest fitness score was selected as the starting configuration for molecular dynamics (MD) simulation.

All-atom Molecular Dynamics Simulation of the protein-ligand complex has been performed using the AMBER 22 software package [24]. GAFF2 (General Amber Force Field 2) has been used to obtain the force-field parameters for the ligand. The ligand’s atomic partial charges was obtained using the Restrained Electrostatic Potential (RESP) technique implemented in the PyRED server. The AMBER ff19SB force field was used for the protein. The required number of counterions (Na^+^) were added to neutralize the whole system. The system was solvated in a truncated octahedron box of the TIP3P water model. The energy minimization was performed for the solvated system. Then the system was heated from 0 to 300 K for 50 ps using Langevin thermostat followed by 50 ps density equilibration and constant pressure equilibration for 500 ps. The final production run was performed for 500 ns at 300 K and 1 bar pressure.

### Constraint Network Analysis

To construct a network of the wild EGFR (with and without Gefitinib), we provided the coordinate obtained from the 500 ns trajectory and a single structure has been used for both the cases **[28]**. All water molecules and buffer ions were removed prior to network construction. This choice is supported by previous work [25, 26], which showed that excluding structural water generally causes negligible changes in protein flexibility indices.

All covalent bonds within EGFR were treated as permanent constraints. Noncovalent interactions, including hydrogen bonds, salt bridges, and hydrophobic contacts, were also incorporated. Hydrogen bond energies (E_HB_) were assigned using an empirical potential [27]. In the holo system, interactions between Gefitinib and surrounding EGFR residues were treated equivalently to protein–protein interactions, allowing analysis of ligand-induced changes in global and local flexibility.

## Results and Discussion

### Quantification of the Global Indices

#### Wild EGFR without Gefitinib

The global flexibility index provides a macroscopic measure of how flexibility and rigidity are distributed within the EGFR constraint network [27]. A commonly used global index is the number of internal independent degrees of freedom (floppy modes, *F*), which reflect the dihedral rotational freedom in the network. Monitoring *F* (or its normalized form, the floppy-mode density *U*) across energy cutoffs allows detection of rigidity–flexibility transitions within the protein. For the wild-type EGFR without Gefitinib, the floppy-mode density (*U*) decreases steadily as the hydrogen-bond energy cutoff (Ecut) increases (**Fig. 1a**). This monotonic decline indicates that the protein becomes progressively more rigid when weaker hydrogen-bond constraints are removed at higher cutoffs. Such a continuous behavior suggests the absence of a pronounced phase-transition like event during the rigidity dilution of EGFR. Thus, the apo EGFR gradually shifts from a more flexible to a more rigid state as E_cut_ increases, reflecting its intrinsic structural stability in the absence of the ligand.

**Fig. 1:**
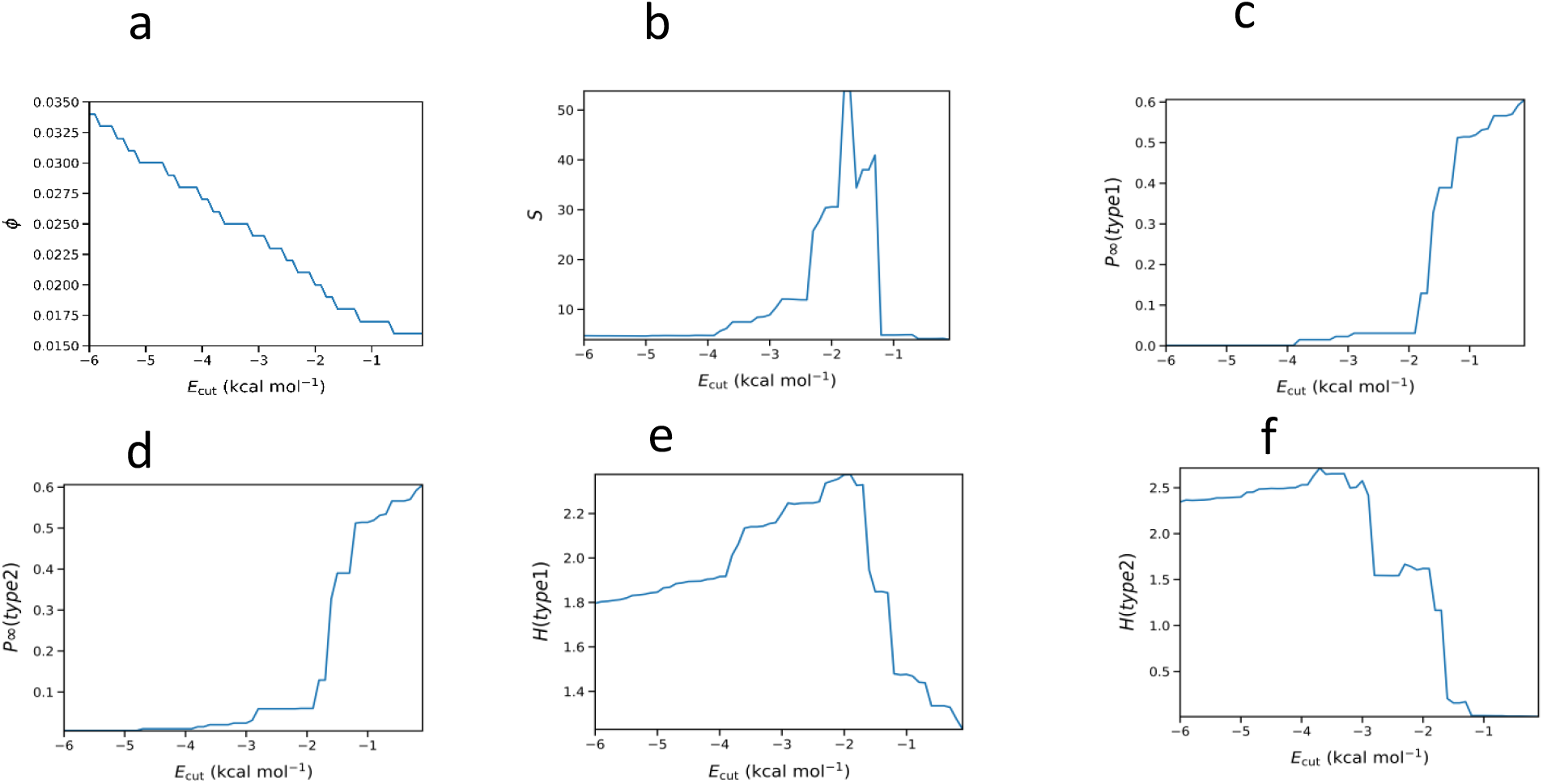
**Representation of the Global Indices for the thermal unfolding simulation of the EGFR (without Gefitinib) as a function of hydrogen bonding energy cutoff E_cut_. a. floppy mode density, b. mean cluster size, c. rigidity order parameter type 1, d. rigidity order parameter type 2, e. cluster configuration entropy type 1, f. cluster configuration entropy type 2.**

The evolution of the mean cluster size, *s*, demonstrates a characteristic multi-stage response of the EGFR constraint network as hydrogen-bond constraints are progressively weakened (**Fig. 1b)**. At low cutoffs (E_cut_ < −4 kcal mol^−1^), *s* remains small and increases gradually, reflecting a predominance of small, disconnected rigid clusters within a largely flexible network. As E_cut_ approaches approximately ∼ −3 kcal mol^−1^, *s* rises more sharply, culminating in a pronounced maximum near E_cut_ ≈ −2 kcal mol^−1^. This peak signifies the transient formation of a large, system-spanning rigid cluster, which is characteristic of a percolation-like transition. Beyond this point, *s* collapses abruptly, indicating that the removal of additional constraints destabilizes the spanning cluster and restores a highly flexible network.

The type-1 rigidity order parameter exhibits behavior consistent with the cluster-size profile. For strongly constrained networks (E_cut_ < −3 kcal mol^−1^), the order parameter remains near zero, indicating the absence of long-range rigidity (**Fig. 1c**) A marked increase occurs at **E**_**cut**_ **≈ −2 kcal mol**^**−1**^, where P∞ rises steeply, signaling the sudden emergence of a macroscopic rigid cluster that spans a substantial portion of the protein. This sharp transition reflects the onset of global mechanical cooperativity. At higher cutoffs, P∞ continues to increase slightly before plateauing, consistent with partial retention of long-range rigidity prior to complete network dissolution.

The type-2 rigidity order parameter follows a similar pattern but exhibits a somewhat more gradual initial rise (**Fig. 1d**) At low E_cut_ values, P_∞_ (type 2) remains negligible, highlighting that rigidity is confined to local regions. A steep transition again occurs at **E**_**cut**_ **≈ −2 kcal mol**^**−1**^, though the increase is smoother than that of type 1, reflecting the metric’s sensitivity to both rigid-cluster extent and internal connectivity. Beyond the transition point, P_∞_ (type 2) increases further, indicating the formation of more cohesive rigid domains before ultimately decreasing as the network becomes fully flexible.

The type-1 cluster configuration entropy provides a measure of the diversity of local cluster arrangements (**Fig. 1e**) For apo EGFR, H (type 1) increases progressively as E_cut_ rises from −6 to approximately −3 kcal mol^−1^, indicating a growing number of possible ways to distribute constraints among small rigid clusters. A maximum is observed near **E**_**cut**_ **≈ −2 kcal mol**^**−1**^, corresponding to a highly heterogeneous network containing an optimal mixture of rigid and flexible regions. Following this maximum, H (type 1) declines sharply, reflecting a loss of configurational diversity as the network abruptly organizes into a more uniform, system-spanning rigid structure. This behavior is characteristic of a second-order–like transition in rigidity analysis.

In contrast, the type-2 cluster configuration entropy remains comparatively high and stable over a broad range of cutoffs until the system approaches the rigidity transition (**Fig. 1f**).This plateau indicates that, despite local reorganization, the global cluster-diversity landscape remains broad. A pronounced drop occurs at **E**_**cut**_ **≈ −2 kcal mol**^**−1**^, at which point the spanning rigid cluster forms and the number of permissible cluster partitions collapses. This entropy loss signifies a strong reduction in structural degeneracy and reflects a network that becomes increasingly dominated by a single rigid topology.

Taken together, all indices consistently identify E_cut_ ≈ −2 kcal mol^−1^ as the primary rigidity-transition point for wild-type EGFR in its apo form. This transition marks the shift from a predominantly flexible, fragmented constraint network to one that transiently supports a system-spanning rigid cluster before ultimately transitioning to a fully flexible state at higher cutoffs. These results highlight the intrinsic mechanical organisation of EGFR in the absence of ligand binding and provide a quantitative framework for comparing its rigidity properties with those of the ligand-bound system.

### Wild EGFR-Gefitinib complex

The floppy mode density decreases almost linearly with increasing E_cut_, indicating a continuous reduction in the number of internal degrees of freedom as hydrogen-bond constraints are progressively weakened (**Fig. 2a**) Compared to the apo system, the ligand-bound EGFR exhibits a slightly lower Φ across the entire cutoff range, consistent with a more constrained and rigidified network upon Gefitinib binding. The smooth nature of the decline suggests that the ligand stabilizes local structural motifs and prevents abrupt fluctuations in network flexibility.

**Fig. 2:**
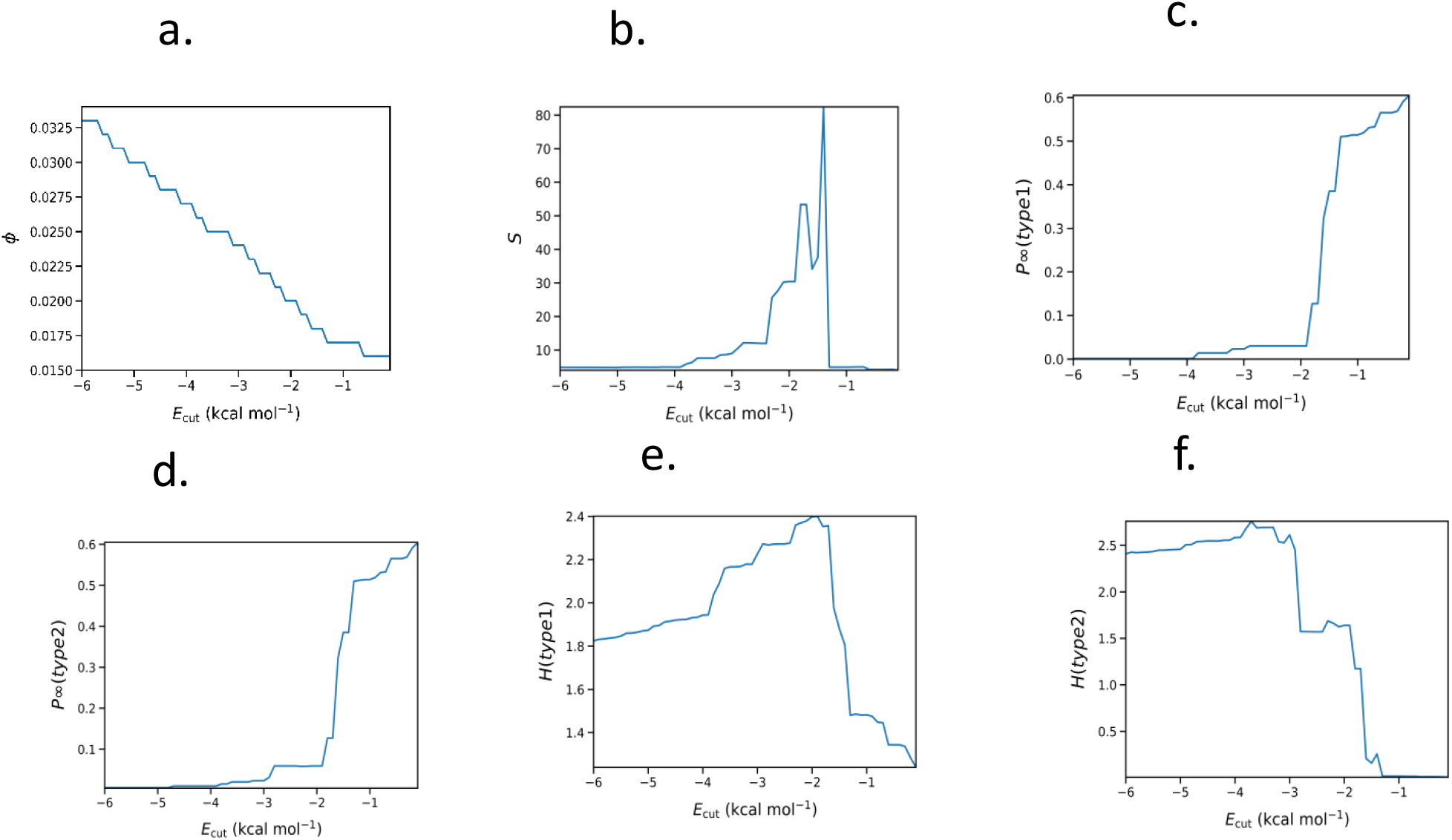
**Representation of the Global Indices for the thermal unfolding simulation of the EGFR-Gefitinib complex as a function of hydrogen bonding energy cutoff E_cut_. a. floppy mode density, b. mean cluster size, c. rigidity order parameter type 1, d. rigidity order parameter type 2, e. cluster configuration entropy type 1, f. cluster configuration entropy type 2.**

**Fig. 3:**
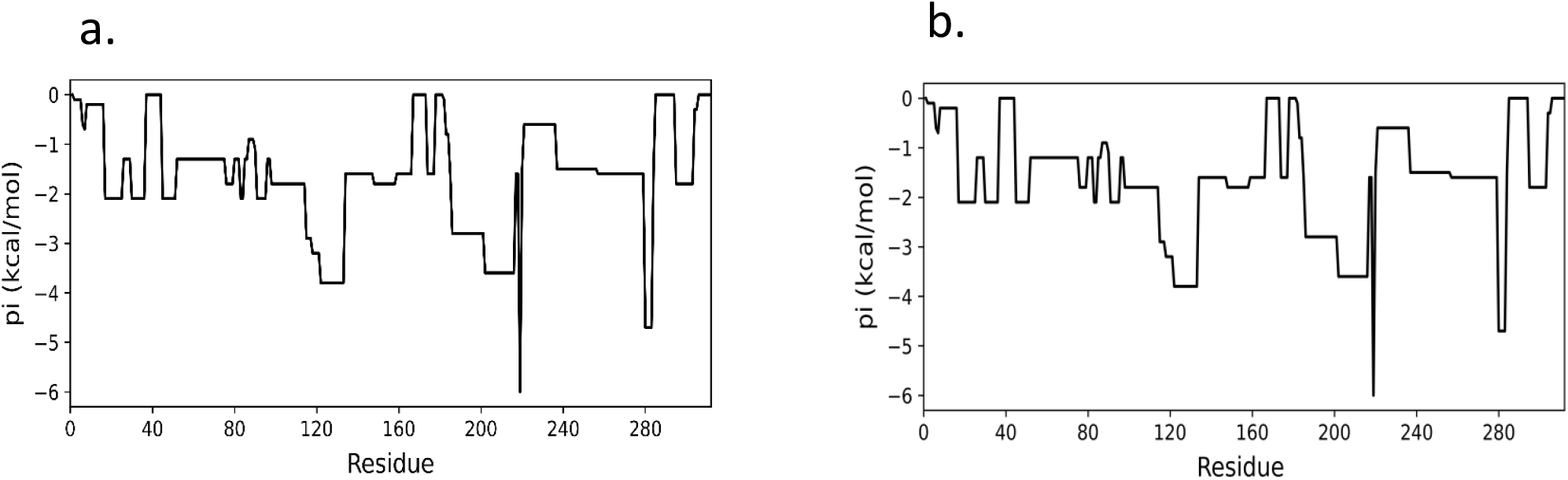
**Quantification of the Percolation index profile for the wild-type EGFR (a) with Gefitinib (holo) and (b) without Gefitinib (apo). The percolation index p_i_ (kcal/mol) reports the local contribution of each residue to network percolation and the formation of a spanning rigid cluster. Negative p_i_ values reflect residues that participate more strongly in the percolating rigid network.**

The mean cluster size shows a prominent peak near **E**_**cut**_**≈ –1.8 kcal mol**^**−1**^, indicating the formation of an extensive, system-spanning rigid cluster (**Fig. 2b**) Notably, the magnitude of this peak is substantially larger than that of the apo state, implying significantly enhanced long-range mechanical coupling in the presence of Gefitinib. Prior to the transition, the gradual increase in *s* reflects incremental growth of medium-sized rigid regions. Following the peak, *s* collapses sharply, signifying the destabilization of the spanning cluster as the network loses critical hydrogen-bond constraints. This marked rise and collapse behavior is characteristic of a strong rigidity percolation transition.

The type-1 rigidity order parameter exhibits a sharp, sigmoidal transition centered at **E**_**cut**_ **≈ – 1.8 kcal mol**^**−1**^, where P∞ increases from nearly zero to ∼0.6 within a narrow cutoff window **(Fig. 2c**). This abrupt jump reflects the rapid emergence of global rigidity in the complex and demonstrates the strong cooperativity introduced by ligand binding. The elevated plateau value of P∞, compared with the apo system, confirms that Gefitinib enhances the extent of long-range mechanical connectivity within EGFR.

A similar transition is observed in the type-2 rigidity order parameter, although P∞ (type 2) displays slightly more gradual growth prior to the main transition (**Fig. 2d**).The steep rise occurring near **E**_**cut**_ **≈ –1.8 kcal mol**^**−1**^ highlights the sudden appearance of cohesive rigid domains that extend across a large fraction of the protein. The higher post-transition value observed for the ligand-bound system indicates that Gefitinib not only triggers the formation of a spanning rigid cluster but also reinforces its internal connectivity and rigidity.

The type-1 cluster configuration entropy initially increases with E_cut_, reaching a maximum at **∼ –1.8 kcal mol**^**−1**^, corresponding to the point of highest structural heterogeneity in the constraint network (**Fig. 2e**). This peak reflects a balance between flexible and rigid regions just before the formation of the spanning rigid cluster. After the transition, H(type 1) decreases sharply, indicating a significant reduction in configurational diversity as the network reorganizes into a more uniform rigid structure. Compared with the apo state, the entropy drop is steeper, suggesting that Gefitinib sharpens the rigidity transition by suppressing intermediate states.

The type-2 cluster configuration entropy remains relatively high and stable at lower cutoffs, reflecting the presence of multiple permissible cluster partitions within the network (**Fig. 2f**). At **E**_**cut**_ **≈ –1.8 kcal mol**^**−1**^, H (type 2) undergoes a pronounced collapse, signaling a dramatic loss of structural degeneracy as the network transitions into a ligand-stabilized rigid topology. The sharper decline and lower final entropy value relative to the apo system imply that Gefitinib significantly restricts the configurational landscape of EGFR, promoting a more ordered and mechanically unified state.

Together, all six global indices consistently identify **E**_**cut**_ **≈ –1.8 kcal mol**^**−1**^ as the primary rigidity transition point for the EGFR–Gefitinib complex. Relative to the apo state, the ligand-bound system displays stronger long-range rigidity, sharper transitions, higher order parameters, and lower post-transition entropy, highlighting the role of Gefitinib in stabilizing the mechanical architecture of EGFR [27].

### Quantification of the Local Indices

#### Comparative Analysis of Percolation index in Wild EGFR and EGFR-Gefitinib Complex

The percolation index (p_i_) provides a quantitative measure of the structural flexibility of individual residues within a protein network [27]. A lower value for a residue indicates that it is part of a larger, more persistent rigid cluster throughout the molecular dynamics simulation, suggesting a reduced local flexibility and increased stability. Conversely, a higher value suggests that the residue is more frequently part of flexible, transient clusters, indicating higher local flexibility. This analysis is instrumental in identifying regions of a protein that undergo significant changes in rigidity upon ligand binding, which can be directly correlated with the mechanism of drug action.

Comparative analysis of the percolation index (p_i_) for wild-type EGFR and the EGFR-Gefitinib complex reveals a striking conservation of the overall profile. The p_i_ values exhibit an almost identical range ∼ 0 to -6 kcal /mol and pattern across the residue span. This high degree of similarity indicates that the binding of Gefitinib does not fundamentally restructure the local network topology of the EGFR kinase domain. The inhibitor’s mechanism, consistent with its ATP-competitive nature, appears to rely on localized steric hindrance and stabilization of a specific inactive state, rather than propagating substantial alterations through the protein’s internal communication pathways. The presence of the inhibitor does not compromise the protein’s highly coupled structural core or significantly re-route its essential network connections.

#### Comparative Analysis of Rigidity Indices in Wild EGFR and EGFR-Gefitinib Complex

This analysis examines the rigidity profile, represented by the rigidity index (r_i_), of the wild-type Epidermal Growth Factor Receptor (EGFR) and the EGFR complexed with the inhibitor Gefitinib (EGFR-Gefitinib complex) [27]. The rigidity index plots (Fig. 4) illustrate the residue-specific energetic contributions to the stability and flexibility of the protein structure. A more negative r_i_ value generally indicates a more rigid residue or region, while values closer to zero indicate more flexible or less stable regions.

**Fig. 4:**
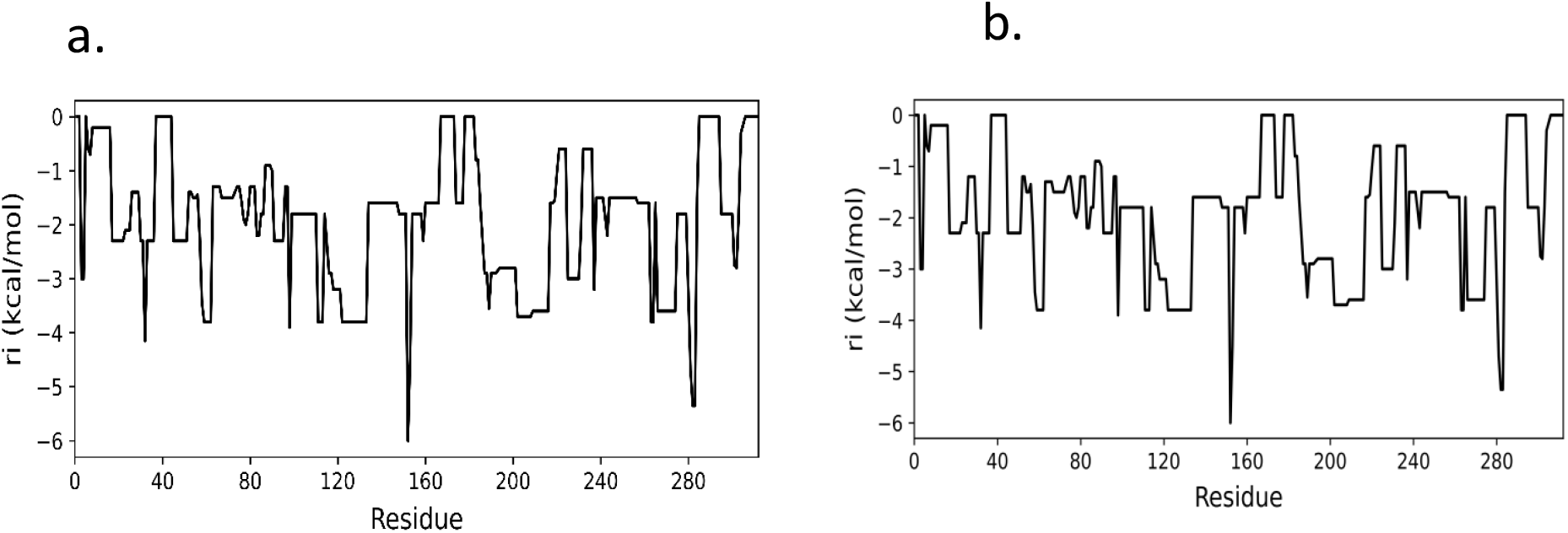
**Quantification of the Rigidity index profile for the wild-type EGFR (a) with Gefitinib (holo) and (b) without Gefitinib (apo).**

The comparative analysis of the rigidity index plots for wild-type EGFR and the EGFR-Gefitinib complex reveals a high degree of conservation in the global energetic rigidity landscape between the two states, suggesting that Gefitinib binding does not induce a dramatic alteration in the protein’s overall fold or stability profile. Both proteins exhibit r_i_ values predominantly ranging from 0 to -6 kcal/mol, with highly rigid core regions and distinct flexible loops maintained across both systems. However, a residue-specific examination is crucial, as the inhibitory action of Gefitinib is postulated to arise from localized stabilization of the inactive conformation within the ATP-binding pocket. These subtle, localized shifts— particularly a trend toward increased rigidity i.e, more negative r_i_ in key catalytic residues— are sufficient to kinetically constrain the dynamic motions necessary for ATP binding, turnover, and subsequent signal transduction, thus explaining the observed therapeutic efficacy despite the largely conserved global rigidity profile.

## Conclusion

In this work, Constraint Network Analysis was employed to systematically characterize the global and local rigidity–flexibility architecture of wild-type EGFR in its apo form and in complex with the ATP-competitive inhibitor gefitinib. Analysis of multiple global rigidity indices reveals that apo EGFR undergoes a well-defined rigidity percolation transition at E_cut_ ≈ −2 kcal mol^−1^, reflecting an intrinsically stable mechanical framework that supports transient long-range rigidity without abrupt structural collapse. Upon gefitinib binding, this transition shifts to a slightly higher cutoff (E_cut_ ≈ −1.8 kcal mol^−1^) and becomes markedly sharper, accompanied by enhanced long-range mechanical coupling, higher rigidity order parameters, and reduced configurational entropy. These features indicate ligand-induced stabilization of the EGFR mechanical network rather than a qualitative reorganization.

Residue-level analyses further demonstrate a striking conservation of both percolation and rigidity index profiles between the apo and ligand-bound states. The absence of large-scale changes in local rigidity patterns suggests that gefitinib binding does not disrupt the intrinsic mechanical topology of the kinase domain. Instead, inhibition appears to arise from subtle, localized rigidification within functionally critical regions, consistent with stabilization of an inactive conformational ensemble rather than global mechanical remodeling.

Collectively, these findings establish wild-type EGFR as a mechanically robust kinase whose rigidity–flexibility landscape is largely invariant to ligand binding. This work provides the first CNA-based mechanical reference state for EGFR and highlights the utility of network rigidity analysis in resolving functionally relevant, yet structurally subtle, effects of small-molecule inhibitors. The framework presented here offers a quantitative baseline for future investigations of oncogenic mutations and drug-resistant EGFR variants, where perturbations to the mechanical network may underlie altered inhibitor sensitivity.

## Acknowledgment

K.B. is thankful to Ashoka University for the doctoral fellowship. K.B. also acknowledges the Indian Council of Medical Research (ICMR) (BMI/11(92)/2022) for providing the SRF research fellowship. We are thankful to the HPC, Ashoka University for providing the necessary infrastructure to help us successfully conduct our research work. The authors sincerely thank Dr. Sapna Sarupria for suggesting the use of Constraint Network Analysis (CNA) in this work.

## Data and Software Availability

All data supporting the conclusions of this work, including MD derived coordinate files and CNA-derived global and local indices, are provided as a compressed ZIP archive (**SI_MD_CNA_EGFR.zip**).

Molecular docking was performed using GOLD (version 2020.3).

AMBER22 software package was used to perform Molecular Dynamics Simulation.

Constraint Network Analysis was performed using CNAnalysis web interface (http://www.cnanalysis.de).

All software used in this study is commercially available or freely available for academic use.

## References

1. K. Henzler-Wildman and D. Kern, Nature, 2007, 450, 964–972.

2. T. J. Kamerzell and C. R. Middaugh, J. Pharm. Sci., 2008, 97, 3494–3517.

3. C. Pfleger, S. Radestock, E. Schmidt and H. Gohlke, J. Comput. Chem., 2013, 34, 220–233.

4. K. Opron, K. Xia and G.-W. Wei, J. Chem. Phys., 2014, 140, 234105.

5. C. Forrey, J. F. Douglas and M. K. Gilson, Soft Matter, 2012, 8, 6385–6392.

6. M. E. Gasper and P. Csermely, Brief. Funct. Genomics, 2012, 11, 443–456.

7. T. B. Mamonova, A. V. Glyakina, O. V. Galzitskaya and M. G. Kurnikova, Biochim. Biophys. Acta, Proteins Proteomics, 2013, 1834, 854–866.

8. Z. Shahbazi and A. Demirtas, J. Comput. Inf. Sci. Eng., 2015, 15, 031009.

9. S. W. Englander and N. R. Kallenbach, Q. Rev. Biophys., 1983, 16, 521–655.

10. J. C. Smith, Q. Rev. Biophys., 1991, 24, 227–291.

11. A. G. Palmer III, Chem. Rev., 2004, 104, 3623–3640.

12. K. Bhattacharjee and A. Ghosh, PLoS One, 2025, 20, e0313738.

13. K. Bhattacharjee, A. Sengupta, R. Kumar and A. Ghosh, Front. Bioinform., 2025, 5, 1536783.

14. D. J. Jacobs, A. J. Rader, L. A. Kuhn and M. F. Thorpe, Proteins, 2001, 44, 150–165.

15. D. J. Jacobs and M. F. Thorpe, Phys. Rev. Lett., 1995, 75, 4051–4054.

16. M. F. Thorpe, M. Lei, A. J. Rader, D. J. Jacobs and L. A. Kuhn, J. Mol. Graphics Modell., 2001, 19, 60–69.

17. D. M. Krüger, P. C. Rathi, C. Pfleger and H. Gohlke, Nucleic Acids Res., 2013, 41, W340–W348.

18. S. Radestock and H. Gohlke, Proteins, 2011, 79, 1089–1108.

19. A. J. Rader and S. M. Brown, Mol. BioSyst., 2011, 7, 464–471.

20. S. Dasetty, J. W. P. Zajac and S. Sarupria, Mol. Syst. Des. Eng., 2023, 8, 1355–1370.

21. Y. Shan, A. Arkhipov, E. T. Kim, A. C. Pan and D. E. Shaw, Proc. Natl. Acad. Sci. U.S.A., 2013, 110, 7270–7275.

22. M. Huse and J. Kuriyan, Cell, 2002, 109, 275–282.

23. G. Jones, P. Willett, R. C. Glen, A. R. Leach and R. Taylor, J. Mol. Biol., 1997, 267, 727–748.

24. D. A. Case, H. M. Aktulga, K. Belfon et al., AMBER 2022, University of California, San Francisco, 2022.

25. H. Gohlke, L. A. Kuhn and D. A. Case, Proteins, 2004, 56, 322–337.

26. T. Mamonova, B. Hespenheide, R. Straub, M. F. Thorpe and M. Kurnikova, Phys. Biol., 2005, 2, S137–S147.

27. B. I. Dahiyat, D. B. Gordon and S. L. Mayo, Protein Sci., 1997, 6, 1333–1337.

28. Supplementary Information: “SI_MD_CNA_EGFR.zip”.

